# Quantification of HER2-low and ultra-low expression in breast cancer specimens by quantitative IHC and artificial intelligence

**DOI:** 10.1101/2025.01.22.634230

**Authors:** Frederik Aidt, Elad Arbel, Itay Remer, Oded Ben-David, Amir Ben-Dor, Daniela Rabkin, Kirsten Hoff, Karin Salomon, Sarit Aviel-Ronen, Gitte Nielsen, Jens Mollerup, Lars Jacobsen, Anya Tsalenko

## Abstract

Recent results of clinical trials in antibody drug conjugate therapies have significantly broadened treatment options for the HER2 low and ultra-low breast cancer patients. However, sensitive, accurate and quantitative evaluation of HER2 expression based on current immunohistochemistry (IHC) assays remain challenging, especially in low and ultra-low HER2 expression ranges.

We developed a novel methodology for quantifying HER2 protein expression, targeting breast cancer cases in the HER2 IHC 0 and 1+ categories. We measured HER2 expression using quantitative IHC (qIHC) that enables precise and tunable HER2 detection across different expression levels as demonstrated in formalin-fixed paraffin-embedded cell lines. Additionally, we developed an AI-based interpretation of HercepTest™ mAb pharmDx (Dako Omnis) (HercepTest™ mAb) using qIHC measurements as the ground truth. Both methodologies allowed spatial resolution and visualization of low and ultra-low levels of HER2 expression across entire tissue sections to demonstrate and enable quantification of heterogeneity of HER2 expression.

Serial sections of 82 formalin-fixed paraffin-embedded tissue blocks of invasive breast carcinoma with HER2 IHC scores 0 or 1+ were stained with H&E, HercepTest™ (mAb), qIHC and p63, then scanned and digitally aligned. Tumor areas were manually selected and reviewed by expert pathologists. HER2 expression was quantitatively evaluated based on the qIHC assay in each 128×128µm^2^ area within tumor regions. We observed statistically significant differences in HER2 expression between IHC 0, 0<IHC<1+, and IHC 1+ groups, and high degree of spatial heterogeneity of the HER2 expression levels within the same tissue, up to five-fold in some cases. We demonstrated high slide-level tumor region agreement of estimates of HER2 expression between the AI-based interpretation of HercepTest™ mAb and the qIHC ground truth with a Pearson correlation of 0.94, and R^2^ of 0.87.

The developed methodologies can be used to stratify HER2 low-expression patient groups, potentially improving the interpretation of IHC assays and maximizing therapeutic benefits. This method can be implemented in histology labs without requiring a specialized workflow.

## Introduction

The evaluation of HER2 in breast cancer has evolved substantially since the 1998 FDA approval of trastuzumab (Herceptin®), alongside the introduction of immunohistochemistry (IHC) assays to guide patient selection.^1,2^ These assays were developed to distinguish HER2-positive tumors (IHC 3+ or IHC 2+ with gene amplification), eligible for trastuzumab treatment, from HER2-negative tumors (IHC 0, IHC 1+ or IHC 2+ without gene amplification), which were not considered for therapy.^1^ This classification enabled effective identification of patients most likely to benefit from HER2-targeted treatment.

Recently, the emergence of antibody-drug conjugates (ADCs) such as trastuzumab deruxtecan (T-DXd; Enhertu®) has expanded therapeutic options to include HER2-low breast cancer, defined as IHC 1+ or IHC 2+ without amplification, based on results from the DESTINY-Breast04 trial (NCT03734029).^3,4^ Furthermore, early findings from the ongoing DESTINY-Breast06 trial (NCT04494425) suggest potential benefit for patients with HER2 ultra-low expression, a subset of IHC 0 cases showing incomplete and faint membrane staining in ≤10% of tumor cells as depicted in Figure 1.^3,5^ As a result, low and ultra-low HER2 expression have gained new clinical relevance.^4^ However, since traditional IHC assays were originally designed to identify highly expressed HER2-positive cases, they are not optimal for reliably distinguishing these lower-expression categories. This challenge is further compounded by high interobserver variability,^6–10^ a problem that was partly solved by standardizing training.^11^ The higher sensitivity of the recently introduced HercepTest™ mAb pharmDx (Dako Omnis) assay may allow detection of more patients with HER2-low and ultra-low expression and enable the investigation of clinical response rate and outcome in these ranges.^12–14^ However, two significant challenges still remain in adapting existing diagnostic tools to these newly actionable subgroups, the adaptation of HER2 detection and interpretation methods.

**Figure 1.**
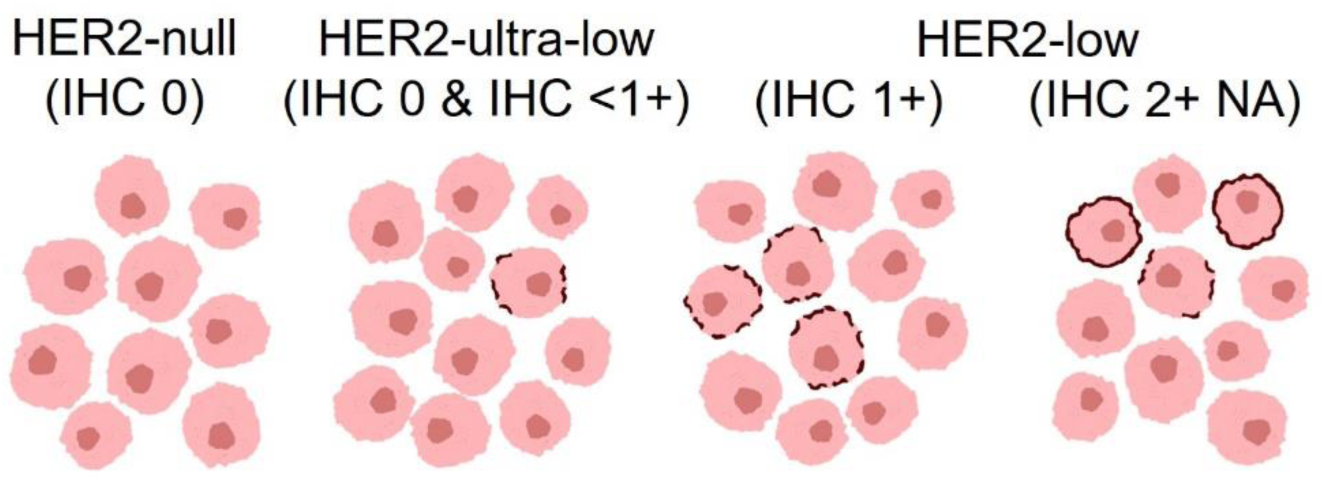
Definitions of HER2 negative tumors used in this study: HER2-null with no membrane staining; HER2-ultra-low with weak membrane staining in less than 10% of tumor cells; and HER2-low with very weak to moderate membrane staining in more than 10% of tumor cells and non-amplified (NA) *HER2* ISH status. These definitions align with the European Society for Medical Oncology consensus statements presented by Tarantino et al., 2023.^26^

Alternative spatial quantitative assays that address the challenges of traditional HER2 IHC, particularly in accurately detecting low and ultralow levels of HER2 expression have been recently developed. Two such methods are quantitative immunofluorescence (QIF)^15^ and RNAscope^16^, the first enables spatial measurement of HER2 protein by quantifying fluorescence intensity calibrated against mass spectrometry cell-line sample controls, providing a continuous objective quantification range^15^. The second enables detection of HER2 mRNA expression directly in formalin-fixed paraffin-embedded (FFPE) tissue sections^16^, which has been reported to correlate with patient responses to T-DXd treatment. Notably, both QIF and RNAscope currently do not differentiate well between tumors scored as IHC 0 and IHC 1+^2,16^.

An additional method is quantitative IHC (qIHC), which may provide a practical and clinically adaptable approach for determining HER2 expression in FFPE tissue.^17,18^ This sensitive quantitative method generates countable chromogenic dots that correspond to HER2 protein levels in tissue sections or cell lines. Unlike conventional IHC, qIHC minimizes the influence of staining variability, resulting in reproducible and quantitative measurements. The assay has demonstrated a lower limit of detection than traditional IHC assays, making it particularly suited for assessing HER2 expression in the low and ultralow range.

The second challenge addresses the issue of HER2 IHC interpretation, particularly in the low-expression range. AI-based methods provide quantitative objective measures of HER2 IHC expression with potential advantages over the currently used human-based practice.^19–23^ In this approach, AI-based algorithms are applied to digitally scanned whole slide images (WSIs) of HER2 IHC stained tissue sections to provide valuable assistance in the interpretation of breast cancer specimens and reduce inter-observer variability, especially in cases with heterogeneous expression or diagnostically challenging HER2-low tumors.^19–22^ Moreover, deep learning–based analysis of WSIs has been shown to enhance patient stratification, resulting in improved progression-free survival and greater accuracy in identifying eligible patients compared to conventional manual scoring based on ASCO/CAP guidelines.^23^ A central challenge with most of these AI approaches is the need to obtain reliable manual annotations, at the cell, region, or WSI level, to establish ground truth for algorithm development. A common practice is to obtain consensus annotations of multiple pathologists per sample in order to resolve disagreement among annotators. This challenge is exacerbated for HER2-low cases where rater agreement is particularly low.^6–10^ Moreover, a recent study comparing AI models for HER2 scoring showed that agreement among automated models was similar to agreement among pathologists,^24^ potentially due to the models being trained on subjective, manually scored data. Therefore, relying on objective ground truth derived from quantitative assays may offer a more robust and reproducible foundation for AI-based interpretation of IHC assays.^25^

In this study, we introduce a modified, tunable qIHC system designed to improve sensitivity for detecting HER2 expression in the low and ultra-low range. By quantitative analysis of qIHC-stained tissue slides, we assess its ability to distinguish among HER2 low, ultra-low and HER2 null expression levels and use qIHC measures as objective ground truth to develop an AI-based model for interpreting HercepTest™ mAb pharmDx (Dako Omnis) stained WSIs. In addition, we present spatial maps revealing heterogeneity of measured and predicted quantitative HER2 expression. This approach could enable reliable, quantitative, and continuous estimation of HER2 expression in low and ultra-low cases, one that can be adapted for future use in clinical settings.

## Material and Methods

### Tissue and Cell Lines

A total of 151 formalin-fixed, paraffin-embedded (FFPE) breast cancer specimens, encompassing both resections and biopsies were screened for this study. From these, 82 specimens having solid tumor areas and characterized as HER2 IHC 0 or 1+ by ASCO/CAP guidelines were selected for algorithm development. All slides were scored by a certified observer according to ASCO/CAP guidelines, and the score was verified by a second certified observer. Discordant cases were discussed and a final score assigned. Cases without invasive cancer or with very sparse tumor representation were not included. One HER2 IHC 0 sample had small 2+ areas and was excluded from group comparison in Figure 4B. According to the criteria outlined in Figure 1, these breast cancer specimens represent HER2-null (n=13), HER2-ultra-low (n=27), and HER2-low (n=42). Cases without invasive cancer or with very sparse tumor representation were not included. All specimens in the study were residual, de-identified, originated from different individuals, and were procured from commercial sources or local hospitals with no possibility for patient identification. The specimens used came from 12 different sources. The study was performed in agreement with the World Medical Association Declaration of Helsinki (2013) and no study protocol was submitted for Ethics Committee approval as this type of analytical study is exempt from Ethics Committee approval in Denmark. FFPE blocks containing cell lines (CCLs) MDA-MB-468, MDA-MB-231, MDA-MB-175, MDA-MB-453, and SK-BR-3 used in this study were originally described and used in the study by Jensen et al., 2017.^18^

### Sectioning

Sections were cut at 4 µm on a Leica microtome (RM2255, Leica Biosystems, Wetzlar, Germany), mounted on K8020 FLEX IHC Microscope Slides (Agilent Technologies, Glostrup, Denmark) and baked for 1 hour at 60 °C. Six consecutive sections from the breast cancer specimens were cut to ensure spatial proximity between the different staining results (see supplementary Figure S1).

### Staining of Tissue Sections

Staining with H&E (CS700 and CS701, Agilent Technologies, Glostrup, Denmark) was performed according to instructions by the manufacturer on a Dako Coverstainer (Agilent Technologies, Glostrup, Denmark). Slides were stained with HercepTest™ mAb pharmDx (Dako Omnis) (GE001, Agilent Technologies, Glostrup, Denmark) or monoclonal mouse-anti human p63 Protein Clone DAK-p63 (Dako Omnis) (GA662, Agilent Technologies, Glostrup, Denmark), (supplementary Figure S1) according to the manufacturer’s instructions. Following staining, slides were dehydrated in ethanol, transferred to xylene, and then coverslipped on a Sakura TissueTek coverfilm coverslipper (Sakura, Osaka, Japan).

The principle of HER2 qIHC staining that generates countable dots based on the iCARD technology is schematically illustrated in Figure 2A. The staining was essentially performed as previously described^17,18^ on an Autostainer Link 48 (Agilent Technologies, Glostrup, Denmark) with a few modifications. The highly sensitive monoclonal rabbit anti-HER2 antibody (DG44) from HercepTest™ mAb was used as the primary antibody in Step 1 at a concentration of 1.5 µg/mL. A polymer consisting of polyclonal goat anti-rabbit antibodies conjugated to a dextran backbone with multiple horseradish peroxidase (HRP) enzymes and a fixed concentration of 3 nM unlabeled goat anti-rabbit antibodies was used in Step 2. A fluorescein containing substrate reporter ^17^ was precipitated as shown in Step 3, and an HRP-conjugated F(ab’) antibody recognizing the fluorescein substrate in Step 4 was used to generate DAB-based qIHC dots in Step 5. In this study we also varied the concentration of the HRP-conjugated dextran polymer in Step 2 to control the qIHC dot density. Tissue sections were stained with 2 pM or 4 pM (supplementary Figure S1), cell lines were stained with 0.125 pM to 2 pM. For negative control staining, the primary antibody was omitted from the staining process. Slides were then dehydrated and coverslipped as described for IHC staining.

**Figure 2.**
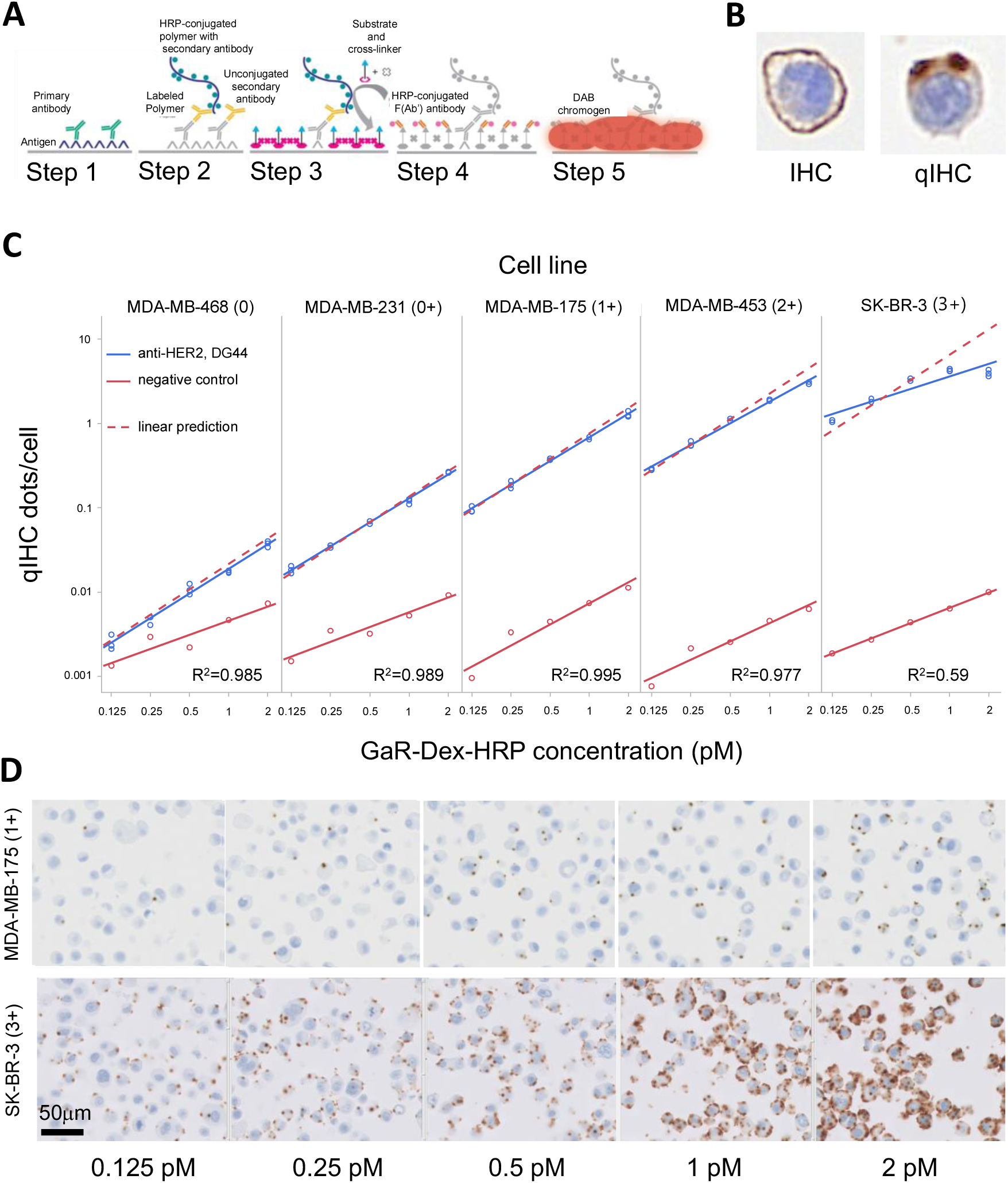
Principle of qIHC. **A**. Schematic illustration of the qIHC staining procedure used in this study, illustration modified from Jensen et al., 2017.^18^ **B.** Example of one MDA-MB-453 FFPE cell stained with standard IHC (HercepTest™ mAb), and one cell stained by HER2 qIHC. **C**. HER2 qIHC dots/cell as a function of the concentration of HRP-conjugated goat anti-rabbit dextran polymer (GaR-Dex-HRP) in five different breast cancer cell lines using the same concentration of the monoclonal rabbit anti-HER2 antibody (clone DG44) tested in triplicates (blue circles, blue linear fit with R^2^ coefficients given) and negative control reagent (antibody diluent without HER2 antibody) in singlets (red circles, red linear fit). The dotted line represents predicted HER2 qIHC dots/cell if doubling or halving the GaR-Dex-HRP concentration calculated from the mean HER2 qIHC dots/cell at 0.5 pM GaR-Dex-HRP. **D.** HER2 qIHC staining at 40X magnification in the MDA-MB-175 (1+) and SK-BR-3 (3+) cell line at different GaR-Dex-HRP concentrations.

### Whole Slide Image Scanning

Scanning was performed on a brightfield Ultra Fast Scanner (Philips, Eindhoven, The Netherlands) at a resolution of 0.25 µm/pixel. WSI images were exported from the Philips IMS in Philips TIFF format (quality 80) for further processing. Additionally, qIHC slides for dot detection model training were scanned on an Axioscan Z.1 with Zen 3.4 acquisition software (Carl Zeiss, Oberkochen, Germany) at a resolution of 0.22 µm/pixel. For each slide, a z-stack of 5 planes was captured with the middle plane being the focus depth and with 1 µm intervals between planes to cover the entire tissue thickness. Extended depth of focus (EDF) using the contrast method in the acquisition software was used to flatten focus in the 5 planes to a single plane.

### Alignment of Whole Slide Image Scans

All WSI scans (Supplementary Figure S1) were aligned to the HercepTest™ mAb WSI scan of the same specimen. Alignment was performed using an in house developed algorithm based on rigid transformations, incorporating only lateral image translation and rotation, to maintain the physical dimensions of tissue sections.

### Tumor Region Annotations on WSI Scans

H&E-stained slides were utilized to identify tumors showing solid growth pattern (solid tumor regions), non-tumor areas, carcinoma *in situ* (CIS), and necrotic areas (Supplementary Figure S2A). Solid tumor regions were included when an estimated ≥ 80% of the cells were invasive tumor. Large areas populated with lymphocytes, stroma, fat cells, normal ducts, necrotic areas, or areas out of focus within the solid tumor regions were excluded. CIS and normal duct exclusion were guided by p63 immunostained sections (Supplementary Figure S2D). Tissue folds that could affect qIHC dot detection were annotated on the qIHC WSI scan for exclusion (Supplementary Figure S2C). Tumor regions were manually annotated on HercepTest™ mAb stained breast carcinoma sections (Supplementary Figure S2B) with the definitions HER2-null, HER2-ultra-low and HER2-low as described in Figure 1, and a HER2 score was assigned based on the ASCO/CAP guideline for the entire specimen.^27,28^ For each case, annotations were done by one trained observer and reviewed by another. Challenging cases were reviewed with a certified breast pathologist to ensure consistency.

### Detection of qIHC Dots

An established UNET AI-model used for nuclei detection and segmentation^29,30^ was adapted to detect qIHC dots by identifying their centroid proximity. The model was iteratively trained using selected fields of view from HER2 qIHC stained WSIs spanning the HER2 IHC-low regime, including HER2-low, ultra-low, and null expression levels. The WSI were initially scanned using Philips Ultra Fast scanner and qIHC dots were manually marked. To enhance accuracy of qIHC dot detection, we used EDF WSIs of the same fields of views scanned on the Zeiss Axioscan Z.1 to improve manual annotation of faint and out-of-focus qIHC dots (Supplementary Figure S3). In addition, qIHC negative controls, i.e., slides stained without the HER2 antibody were used as additional training samples to improve the capability of distinguishing noisy patterns and avoid false-positive dots. Eventually, the AI model used for qIHC dot detection achieved high accuracy with an F1-score of 0.96 in comparison with human annotated ground-truth on the evaluation FOVs.

### qIHC Patch Count Analysis

Patches for training and evaluation of the AI-model were defined around every single nucleus in annotated tumor regions.^29,30^ Nuclei were used as patch centers to ensure preferential sampling of patches that contained cells. Dot concentration at 4 pM HRP-conjugated polymer was chosen to ensure adequate dynamic range in quantification of low HER2 protein expression. Based on this, a patch size of 128 µm × 128 µm, equal to 512 × 512 pixels with a resolution of 0.25 pixel/µm, was used to balance spatial resolution and statistical stability of dot densities. For several slides exhibiting high dot densities, a lower staining concentration at 2 pM of the HRP-conjugated polymer was used to limit dots overlap and ensure accurate quantification. In these cases the dots count was multiplied by a factor of 2 (Supplementary Figure S4). Slide qIHC count for qIHC stained sections was calculated by averaging the patch qIHC dot counts from all patches in annotated tumor areas. For heatmap visualization in Figure 4, patch qIHC counts were computed for patches centered around each pixel in the visualization. For heatmap visualizations in Figure 5, patches were selected on a 32 µm × 32 µm grid, leading to patch overlap of 75%.

### Statistical Analysis

Statistical analyses were performed using the Pingouin version 0.5.4 Python statistical package. The Kruskal-Wallis H-test, a non-parametric one-way ANOVA, was used to compare medians of slide qIHC counts between slides with different overall IHC scores. Where the Kruskal-Walli’s test indicated significance, the Wilcoxon rank-sum post hoc test, with Holm P value adjustment for multiple testing was used to test for significant differences between each pair of overall IHC score groups. Comparison of average slide qIHC counts of HER2-null slides stained with anti-HER2 antibody and negative controls was done using a paired, two-sided t-test with Welch correction for unequal variances (Figure 4B). A single specimen was removed from the qIHC count analysis due to a discrepant HER2 IHC score.

### AI-Model Development for Predicting HER2 Expression

The method for developing AI-models from the standard ResNet50 architecture^31^ included training to quantitatively predict the patch qIHC dot count in the HercepTest™ mAb stained image patches, as illustrated in Figure 5A. The ground truth quantitative HER2 expression for model training was determined by the HER2 qIHC dot count detection on consecutive stained sections and aligned to the HercepTest™ mAb stained section (supplementary Figure S1). To investigate performance of the AI-model development method, k-fold cross-validation analysis was applied splitting the complete specimen dataset into eight non-overlapping subsets, each containing 72 (or 71) training specimens and 10 (or 11) test specimens (see Supplementary Figure S5A). Following the development of a ResNet50 model with training specimens of one subset, this AI-model was then evaluated with the test specimens not used for training of this model. Thereby k-fold cross-validation ensured that variation in all sections was used for testing and proved the overall applicability of the AI-model development method. Within each training subset, tumor regions in each specimen were further divided into non-overlapping training and validation regions creating approximately 80%/20% train-validation non-overlapping image patch samples. During training, patches were sampled to reflect the qIHC dot count distribution within each batch. Additionally, the number of patches sampled from different tissues in the training set was balanced throughout each training epoch to prevent bias towards a specific tissue. Model training optimization was performed using the Adam optimizer and guided by mean-squared-error (MSE) loss, with a reduce-on-plateau learning rate scheduler and early stopping criterion, both informed by validation MSE loss.

## Results

### qIHC quality compared with traditional HER2 IHC

To illustrate the difference of a HER2 qIHC staining versus a traditional HER2 IHC staining, Figure 2B has one example of an FFPE breast cancer cell from the MDA-MB-453 cell line stained by IHC using HercepTest™ mAb and one cell stained by qIHC. In this example the HercepTest™ mAb staining generates a strong DAB staining on the cell membrane, whereas HER2 qIHC staining results in two dots located on the cell membrane, spreading into the cytosol. The dots generated by qIHC are often in the vicinity of the cell membrane, but due to the size of dots and the nature of the dot generating process, they may also locate to other cellular parts as seen here.

### qIHC FPPE-embedded Cell Line Results

To demonstrate the tunability of the qIHC system, different concentrations of the HRP-conjugated polymer were used to generate HER2 qIHC dots in five FFPE breast cancer cell lines with different HER2 expression levels while keeping a constant concentration of the monoclonal rabbit anti-HER2 antibody (clone DG44) (Figure 2C and supplementary Figure S6). The blocks used were the same that were previously characterized by Jensen et al., 2017^18^ These cell lines exhibit HER2 expression levels from 1.5 x 10^3^ to 5.6 x 10^5^ HER2 receptors/cell and have HER2 IHC scores from 0 to 3+^18^, thereby constituting a continuum of membranal HER2 expression. As already demonstrated by orthogonal methods, the qIHC measurements are directly proportional to the HER2 concentration in FFPE cell lines. The measurements in the present study are plotted against the data on ELISA and flow cytometry published by Jensen et al., 2017^18^ to verify the linearity of the qIHC measurements (supplementary Figure S7). Figure 2C illustrates the correlation between number of qIHC dots per cell and the concentration of the HRP-conjugated polymer that amplifies the signal. It is observed that doubling of the polymer concentration leads to doubling of the number of qIHC dots/cell. For the three cell lines MDA-MB-468, MDA-MB-231, and MDA-MB-175 with less HER2 expression, a linear relationship across the tested HRP-conjugated polymer concentration range, from 0.125 pM to 2 pM and number of qIHC dots was seen. At low polymer concentrations, the two cell lines with higher HER2 expression, MDA-MB-453 and SK-BR-3, also had a linear relationship between polymer concentration and qIHC dots/cell, but at the high concentrations of 1 pM and 2 pM this linearity was lost. This is due to a saturating dot density as dots begin to overlap and the dot detection algorithm cannot estimate the actual number of dots. The background of nonspecific dots from the qIHC staining protocol at five different polymer concentrations is also illustrated in Figure 2C, where red dots represent qIHC staining in which the HER2 antibody has been replaced by the negative control reagent at increasing concentrations of the polymer. The mean number of dots per cell is similar across all tested cell lines when using the negative control reagent and increases with increasing polymer concentration. The image patches in Figure 2D upper panel, illustrate how the dot numbers in the MDA-MB-175 cell line double at every concentration increase while remaining within a range where they can be counted without clustering, and thus are within the dynamic range. The dot numbers in the SK-BR-3 cell line is also within the dynamic range at low concentrations, but dots begin to overlap at high polymer concentrations preventing detection of single dots (Figure 2D, lower panel). Conversely, as a significant portion of primary antibodies are now visualized, the staining begins to resemble the traditional IHC membranous staining. Overall, the qIHC results presented in Figure 2 demonstrate the scalability and adaptability of the qIHC method to different expression levels, and how an improved signal-to-noise level can be achieved by adjusting the concentration of the HRP-conjugated polymer.

### qIHC on Breast cancer Specimens

We then applied the qIHC staining method using either the negative control reagent or HRP-conjugated polymer on 82 FFPE breast cancer specimens with consecutive sections stained with H&E, HercepTest™ mAb, and p63 as described in the supplementary Figure S1. Figure 3 shows representative staining results from HER2-low, HER2-ultra-low and HER2-null breast cancer specimens stained with HercepTest™ mAb (left panel) and HER2 qIHC (right panel). The middle panel in Figure 3 represents a digital enhancement to visually clarify the faint/barely perceptible HER2 staining. As anticipated, the qIHC dot number increases from the HER2-null to the HER2-ultra-low and HER2-low in these section patches.

**Figure 3.**
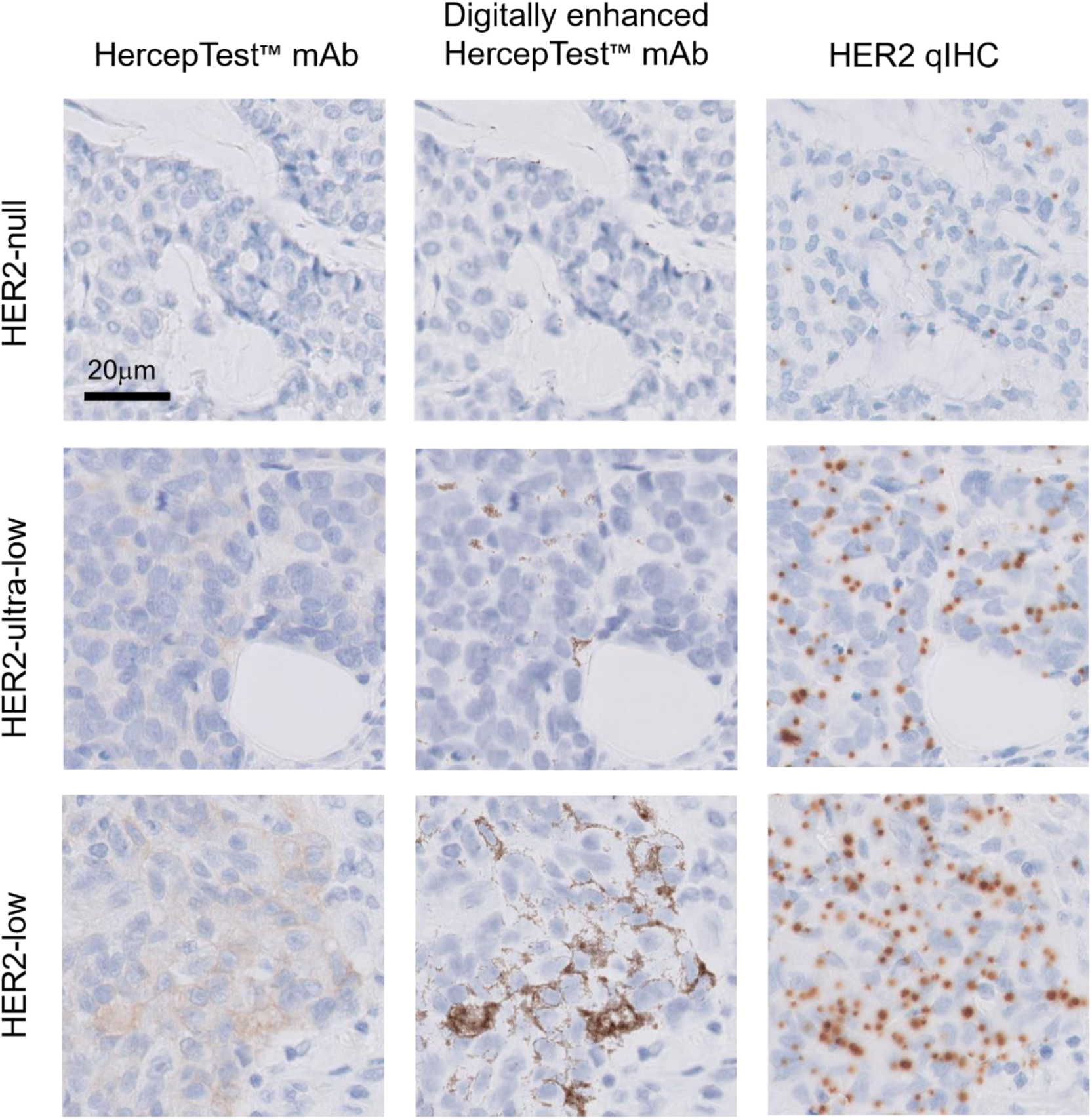
Comparison between IHC, digitally enhanced IHC and qIHC. Top, middle and bottom rows with representative field of views (FOVs) of HER2-null, HER2-ultra-low and HER2-low breast cancer tissues, respectively. Left and right columns have FOVs of aligned sections with HercepTest™ mAb and HER2 qIHC, respectively. The middle column represents the HercepTest™ mAb stained section with a digital enhancement to emphasize faint/barely perceptible membrane staining.

### qIHC Analysis and Spatial Distribution of HER2 Expression

AI-based automated detection of qIHC dots is illustrated in Figure 4A as squared boxes added around recognized dots. We calculated the slide qIHC count for all specimens and illustrated those in the box plot in Figure 4B grouped by HER2 category. The median slide qIHC count is significantly different between the three HER2 categories, and the median number of qIHC dots per image patch is also significantly different when comparing the 13 HER2-null specimens stained with negative control reagent versus staining with the primary anti-HER2 antibody (Figure 4B). For these 13 specimens characterized as HER2-null, the low number of qIHC dots per patch following staining with the negative control reagent illustrates the background level of this system and the high signal-to-noise observed in qIHC. We used a sliding window approach to count dots, allowing us to quantify HER2 expression across the entire tissue sections. In Figure 4C and 4D, we demonstrate such quantitative spatial distribution of HER2 expression from the qIHC dots detected. Figure 4C illustrates a HercepTest™ mAb stained tissue WSI in the HER2 IHC 0 category with manual annotations of solid tumor regions as described in the material and method section and in the supplementary Figure S2. Two small areas with higher magnification are also shown below panel 4C. In Figure 4D a qIHC dots-count heatmap is overlaid on this tissue to reveal the spatial heterogeneity of the quantitative HER2 expression across this WSI, and the same subregions as under Figure 4C is seen under Figure 4D to illustrate this heatmap at higher magnification.

**Figure 4.**
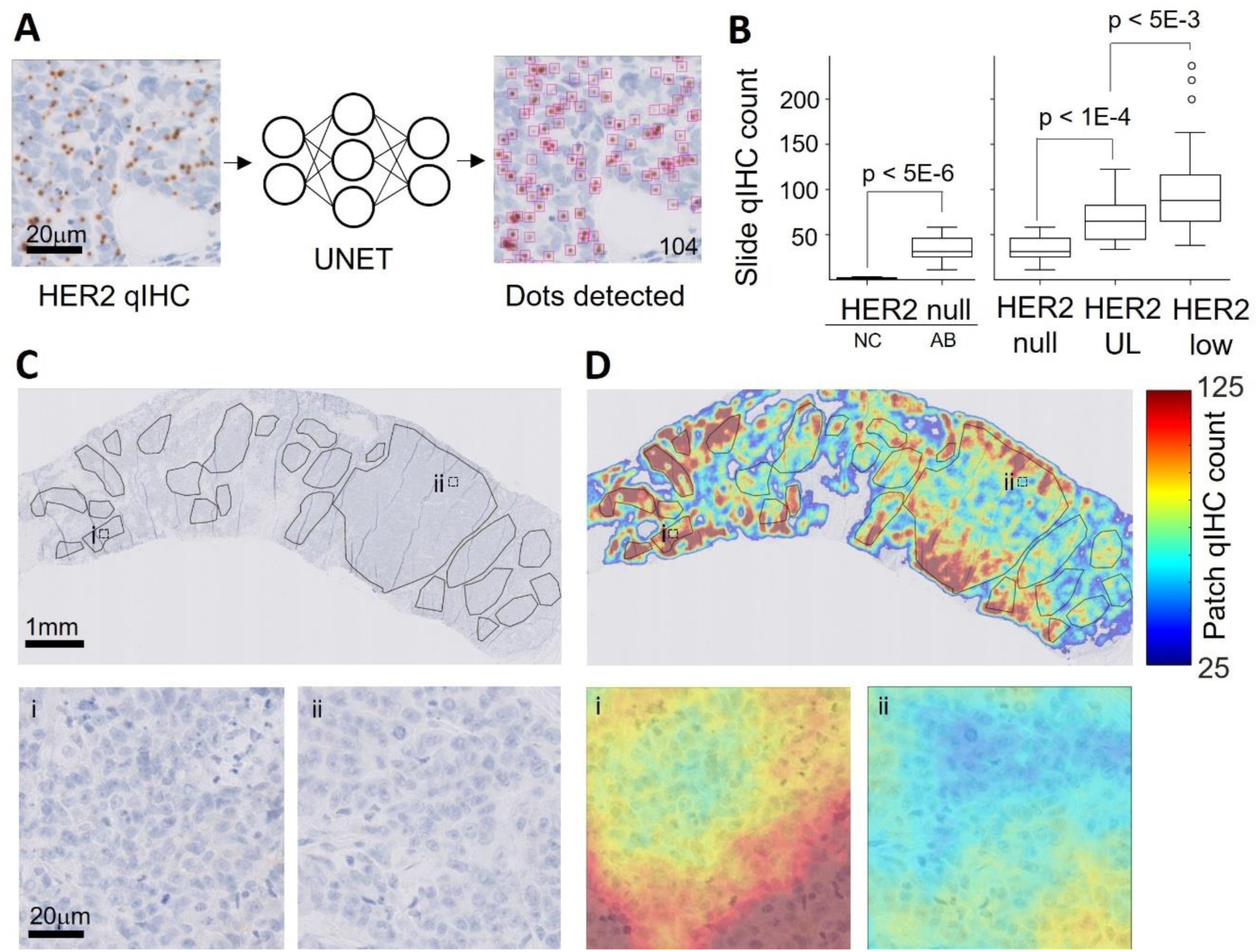
qIHC dot detection and visualization. **A.** Principle of HER2 qIHC dot detection by a UNET deep-neural-network showing detected dots with magenta boxes, and the total number of detected dots in the bottom right corner. **B**. Left panel has the slide qIHC count, calculated by averaging the patch qIHC dot counts from all patches in annotated tumor areas, for HER2-null cases stained with negative control reagents (NC) and HER2 antibody (AB). The p-value is given for a paired, two-sided t-test with Welch correction for unequal variances between the NC and AB groups. Right panel has slide qIHC count in the three categories HER2-null, HER2-ultra-low (UL), and HER2-low (low) and p-values are from median comparison using the Kruskal-Wallis H-test. **C**. Top row has breast cancer tissue WSI stained with HercepTest™ mAb and solid tumor area annotations in black markings. Bottom row has enlarged areas i and ii from top row. **D**. Same specimen as in **C** with the observed patch qIHC count overlaid using a heat map. Blue to red colors indicate low and high patch qIHC count, respectively.

### AI Model Prediction of HER2 Expression Using patch qIHC Count Ground Truth

We then trained AI-models to predict HER2 expression on the HercepTest™ mAb WSI scans using the ground truth HER2 expression from qIHC dot count as described in the Materials and Methods section (Figure 5A). The resulting AI-models could quantitatively predict the number of qIHC dots per patch in WSI of HercepTest™ mAb stained specimens. Figure 5B illustrates the correlation of predicted qIHC dots from such AI-models as a function of the actual number of observed qIHC dots. A Pearson correlation coefficient at 0.94, equal to R^2^ at 0.87 was found. This correlation was found by splitting the entire dataset of 82 specimens into several training and testing subsets as described in the Materials and Methods section, and for each training subset a new AI-model was developed and evaluated on the test subset (Supplementary Figure S5). Using this procedure, we ensured that none of the tested AI-models were trained on the same tissue WSI used for evaluation tests. Figure 5C shows examples of two HER2 expression heatmaps, in which the left one represents the actual HER2 qIHC dots detected, and the right one represents the predicted quantitative HER2 expression output from an AI-model that received the HercepTest™ mAb WSI scan as input. For this specific breast cancer WSI, the Pearson correlation coefficient describing how well the predicted HER2 expression correlated with actual HER2 expression was found at 0.82, with R^2^ at 0.67.

**Figure 5.**
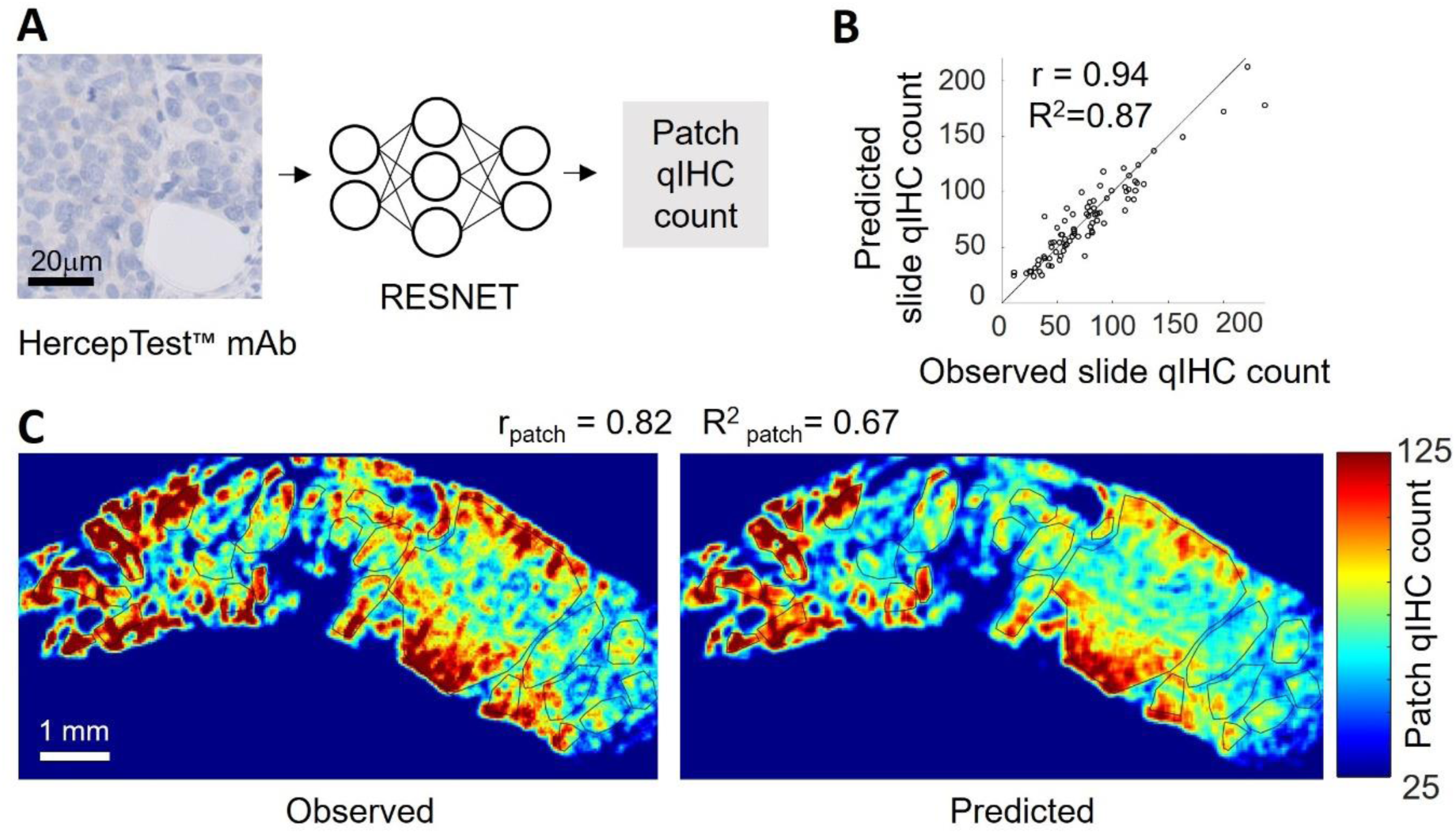
qIHC patch count prediction and visualization. **A.** Principle of patch HER2 qIHC dot prediction by the trained ResNet50 deep neural network. The HercepTest™ mAb WSI scan is processed by the trained ResNet50 AI-model that outputs the predicted numbers of HER2 qIHC dots per patch. **B**. AI predicted HER2 slide qIHC counts as a function of the actual number of HER2 slide qIHC counts calculated from consecutive sections. The correlations given are based on slide comparisons across 82 tissues **C**. Left panel with heatmap from one breast cancer specimen to illustrate patch qIHC dot count of observed HER2 expression, and right panel the AI-predicted HER2 qIHC dots count per patch. Blue to red colors indicate low and high patch qIHC count, respectively. Solid tumor regions from manual annotations are shown with black markings. The correlations given are based on patch comparisons.

## Discussion

In this paper, we introduced a new methodology for quantifying HER2 expression, specifically targeting breast cancer cases in the HER2 low and ultra-low range, using an analysis of qIHC-stained tissue slides. Additionally, we developed an AI-based interpretation of HercepTest™ mAb using qIHC analysis as ground truth.

Recent successes of ADC-based therapies, as demonstrated in DESTINY-Breast04 and DESTINY-Breast06 clinical trials, along with the FDA approval of T-DXd for the treatment of metastatic HER2-positive and HER2-low breast cancer, have expanded the treatment options for the significant subgroup of breast cancer patients with prospects of efficacy for breast cancer patients in the HER2-ultra-low range as well.

However, sensitive, accurate and quantitative evaluation of HER2 expression based on currently approved IHC assays remains challenging, especially in low and ultra-low ranges of HER2.^4^ The ability to correlate and differentiate clinical outcomes with HER2 expression depends not only on the efficacy of the tested drugs but also on the performance of diagnostic assays employed to assess HER2 expression. Advanced computational approaches could improve the interpretation of such IHC assays and could be of high benefit for identifying the best treatment options for current and future HER2-targeted therapies. More importantly, it could provide a basis for objectively assessing the lower level of HER2 protein expression that maximizes therapeutic benefits, while minimizing drug exposure risks for patients. In this study, we utilize the qIHC system to quantify the underlying HER2 expression profile, rather than relying on the semi-quantitative ASCO/CAP score. This qIHC measurement serves as the ground truth for our AI model. While the relationship between qIHC and ASCO/CAP scoring is certainly important, the ASCO/CAP scores are primarily employed in this study to ensure that the various HER2 patient groups are adequately represented. As first suggested by Jensen et al., 2017,^18^ we now demonstrate that qIHC can be used as a tunable system, where the qIHC dot count is directly proportional to the concentration to the dot-generating component. This allows for quantitative HER2 detection with a high signal/noise ratio at different protein expression levels, as demonstrated by our reported reliable and consistent quantitative expression information in FFPE cell lines that have HER2 expression ranges from IHC 0 to 3+. Hence, even though the FFPE tissues used in this study are exclusively HER2 IHC 0 and IHC 1+, the method can be adapted to cover the full HER2 expression range. Additionally, the method can be extended to other cancer types depending on the HER2 pattern of expression as well as intra-tumoral heterogeneity.

We also demonstrate that qIHC slides can be automatically and accurately processed using an AI detection model to identify qIHC dots, enabling scalable processing of qIHC staining. By integrating the detected qIHC dots over patches and averaging them across tumor regions, qIHC can provide continuous spatial quantification of HER2 expression, allowing statistically significant differentiation between various IHC HER2 low categories above the background level of qIHC staining, as shown in Figure 4B. Alternatively, different thresholds could be applied, particularly at ultra-low levels, to provide varying degrees of stratification in future CDx analyses or studies.

In addition, patch-based quantification of HER2 expression using qIHC staining can be spatially mapped across the entire breast cancer tissue, providing valuable insights into HER2 expression heterogeneity. This information can be graphically overlaid on HER2 IHC stained WSI for both visualization and quantification of the heterogeneity, as demonstrated in Figure 4C-D. Furthermore, using qIHC staining, we were able to visualize low levels of HER2 expression that are challenging to detect with the human eye using traditional IHC (subpanels i and ii in Figure 4C-D), or even expression levels below the limit of detection of currently available HER2 IHC assays. While patch-based qIHC scores are currently summarized into global scores, they could conceivably be adapted to offer more detailed and spatially resolved measurements of HER2 expression in low and ultra-low regimes. Additionally, new metrics leveraging qIHC-based spatial analysis of HER2 expression could be developed, which may be particularly useful for studying mechanism and therapeutic effects of Enhertu® or other ADC drugs, such as the bystander effect.

Furthermore, we successfully demonstrated a method for using our qIHC staining and analysis as ground truth for training AI models to extract information from the HercepTest™ mAb pharmDx stain. This method unlocks the ability to infer continuous quantitative expression levels from traditional HER2 IHC. Two key factors contributed to the success of this method. Firstly, the quantitative, high sensitivity, tunability and scalability of qIHC staining and analysis for HER2 expression make qIHC an ideal candidate for providing reliable and consistent ground truth for AI modeling. Secondly, the relatively high sensitivity of HercepTest™ mAb^12^ generates a minute membrane staining signal even in low and ultra-low HER2 expression tissues, although it is difficult to detect by the human eye. This weak signal, which an AI model can learn to detect, is made apparent when the DAB signal is digitally enhanced, as shown in Figure 3. Using our AI model training method, we achieved excellent agreement between model inference and measured qIHC expression at the slide level. This agreement extended to low levels of HER2 expression (Figure 5B). Additionally, strong consistency was observed in the spatial quantification of HER2 expression across the entire breast cancer tissue section, providing valuable insights into HER2 heterogeneity (Figure 5C).

While the results of this study are promising, adding more data and increasing its diversity are important to improve AI model generalizability. In addition, current qIHC analysis faces limitations in differentiating between cytoplasmic and membranous HER2 expression. However, future developments, such as multiplexing membranes markers with qIHC staining using different chromogens and applying algorithmic approaches, could assist in distinguishing between these expression patterns. It should be emphasized that the data provided from this work is purely investigational, and the observations brought forward require additional clinical validation before being employed in the diagnostic space.

Several reports use orthogonal methods as opposed to IHC to examine whether alternative quantitative methods can help improve predictive capabilities while maintaining the important spatial resolution of specimens intact,^16,32^ but these methods require specialized workflows and have so far shown limited resolution in distinguishing HER2 IHC null, HER2 IHC ultra-low and HER2 IHC low groups. As we demonstrate in this paper, a qIHC-based workflow or AI-based interpretation of HercepTest™ mAb could be used to measure the HER2 expression in FFPE specimens without requiring a specialized workflow not currently in place in histology labs, in contrast to other quantitative methods for measuring HER2. Due to the large dynamic range, qIHC technology can be used to separate even low-expression patient groups. Furthermore, qIHC staining is not limited to HER2 but can be extended to any IHC biomarker at any expression level.

Looking ahead, more detailed, automatic, spatial, and continuous quantification of HER2 expression across WSI may provide insights into the impact of HER2 heterogeneity on T-DXd response and help optimize patient selection for treatment decisions. Moreover, the methods presented here could be expanded to cover the full range of HER2 expression and applied to other tumors beyond breast cancer. Additionally, these techniques hold the potential for measuring multiple biomarkers through multiplexing and could even be extended to sparse tumors in the future.

In conclusion, this work illustrates potential for significant advancement in the field of pathology by utilizing the power AI based interpretation of IHC measurements to better quantify and spatially resolve protein expression in cancer tissue, ultimately providing better tools for studying response to therapy.

## Acknowledgements

The authors express their sincere appreciation to Steve Laderman, Lou Welebob, Darlene Solomon, Sam Raha, and Lars Von Gersdorff for their unwavering support and encouragement. Special thanks to Claudia Gottstein, Ghislain Noumsi, Doug Clark, and Emin Orudjev for their invaluable scientific discussions, and to Josh Littrell for statistical assistance. We appreciate the histology assistance provided by Charlotte Aagerup, Anne Falkenskjold, Anne Maarbjerg, and Marianne Marcussen.

## Author Contributions

FA, EA, IR, OB, DR, AB, KH, KS, JM, LJ, and AT contributed to study concept and design. FA optimized IHC assays and generated the data. EA, IR, OB, and DR trained AI models and analyzed the data. AB developed supporting software tools. KH, KS, and SA reviewed and annotated IHC stained tissues. JM, IR, EA, OB, FA, and AT wrote the manuscript, GN edited the manuscript. LJ and AT supervised the research.

## Data Availability Statement

Aggregated data is available on reasonable request.

## Ethics statement and patient consent

All specimens in the study were residual, de-identified, originated from different individuals, and were procured from commercial sources or local hospitals with no possibility for patient identification. The study was performed in agreement with the World Medical Association Declaration of Helsinki (2013). No study protocol was submitted for Ethics Committee approval as this type of analytical study is exempt from Ethics Committee approval in Denmark.

## Supplementary Information

**Figure S1.**
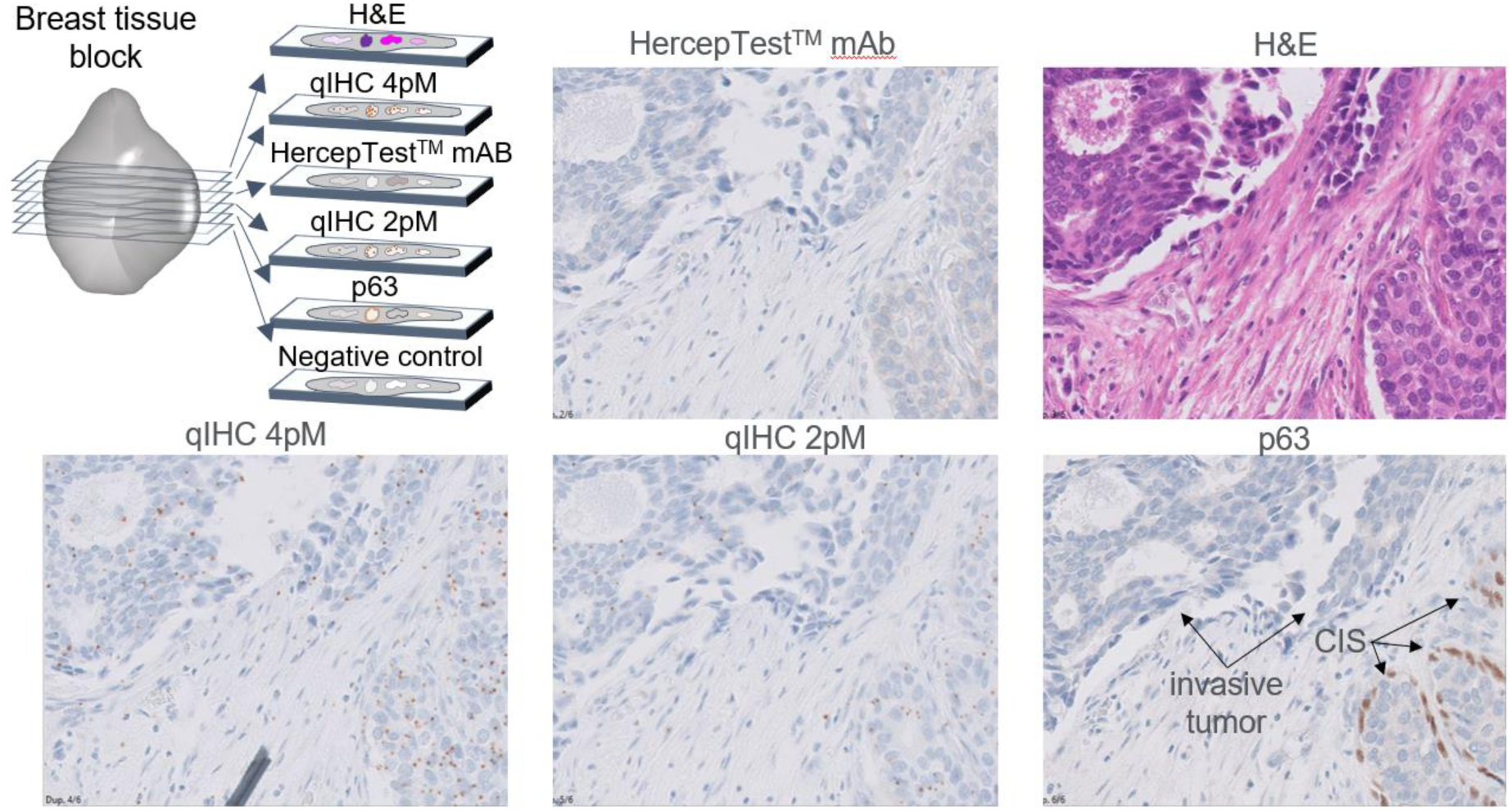
Overview of tissue sections used. Upper, left panel illustrates the six consecutive sections cut from each breast cancer specimen and how these sections were stained. Remaining panels show a representative, aligned field of view for five stained sections with invasive tumor and carcinoma in situ (CIS) indicated in the p63 stained section.

**Figure S2.**
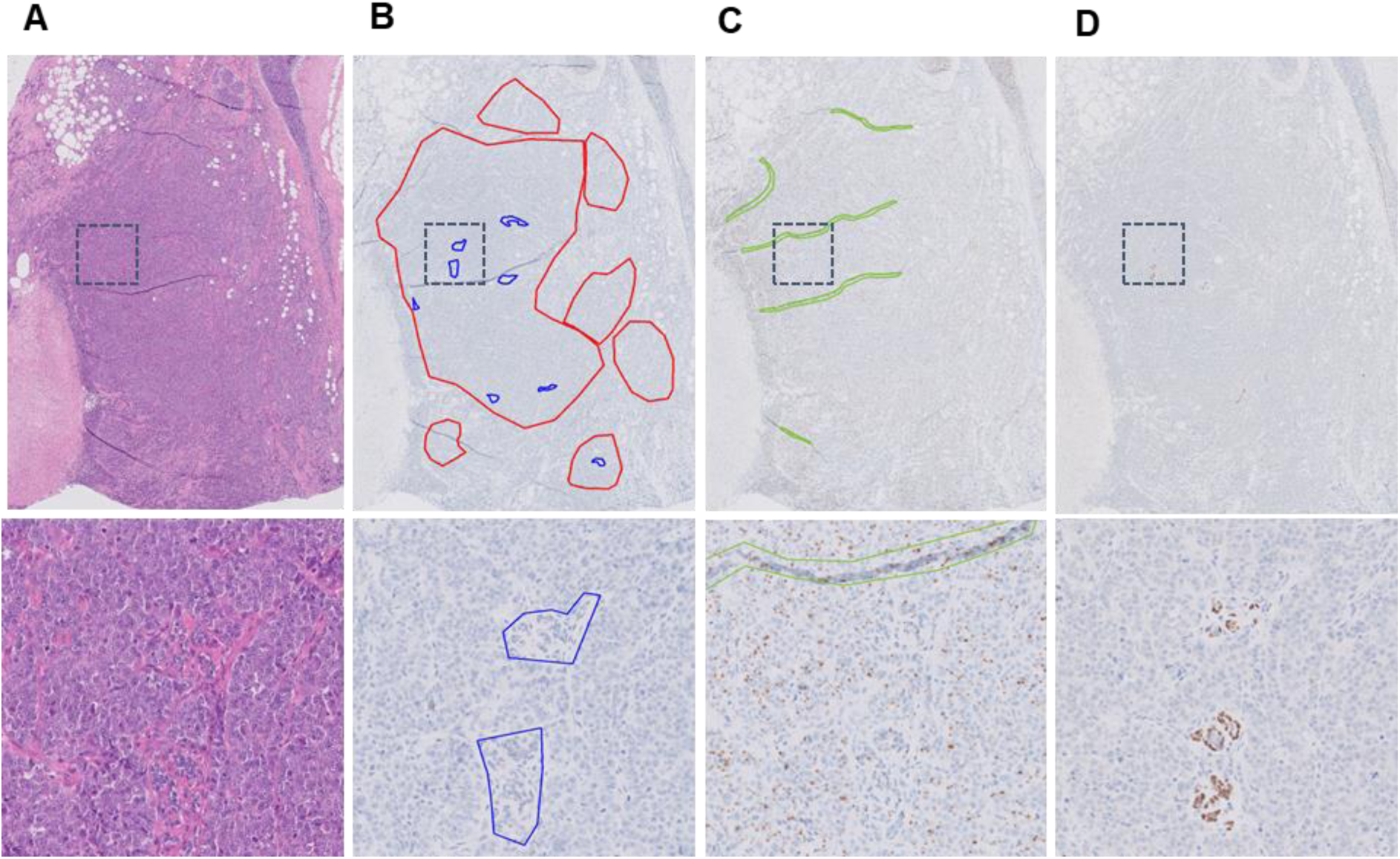
Illustration of tumor annotations in different sections of a breast cancer specimen. Images in the top row represent an overview, and the bottom row a zoomed-in area as indicated by the dashed box. **A**. H&E counterstain to support identification of solid tumor regions and non-tumor areas. **B**. HER2 IHC stain by HercepTest™ mAb where solid tumor regions were encircled red, while large non-tumor areas representing stroma and normal ducts were encircled blue for exclusion. **C**. HER2 qIHC staining for dot detection in which known artefacts such as tissue folds were marked green for exclusion. **D**. The p63 IHC stain supported the identification of normal ducts and CIS structures to be excluded.

**Figure S3.**
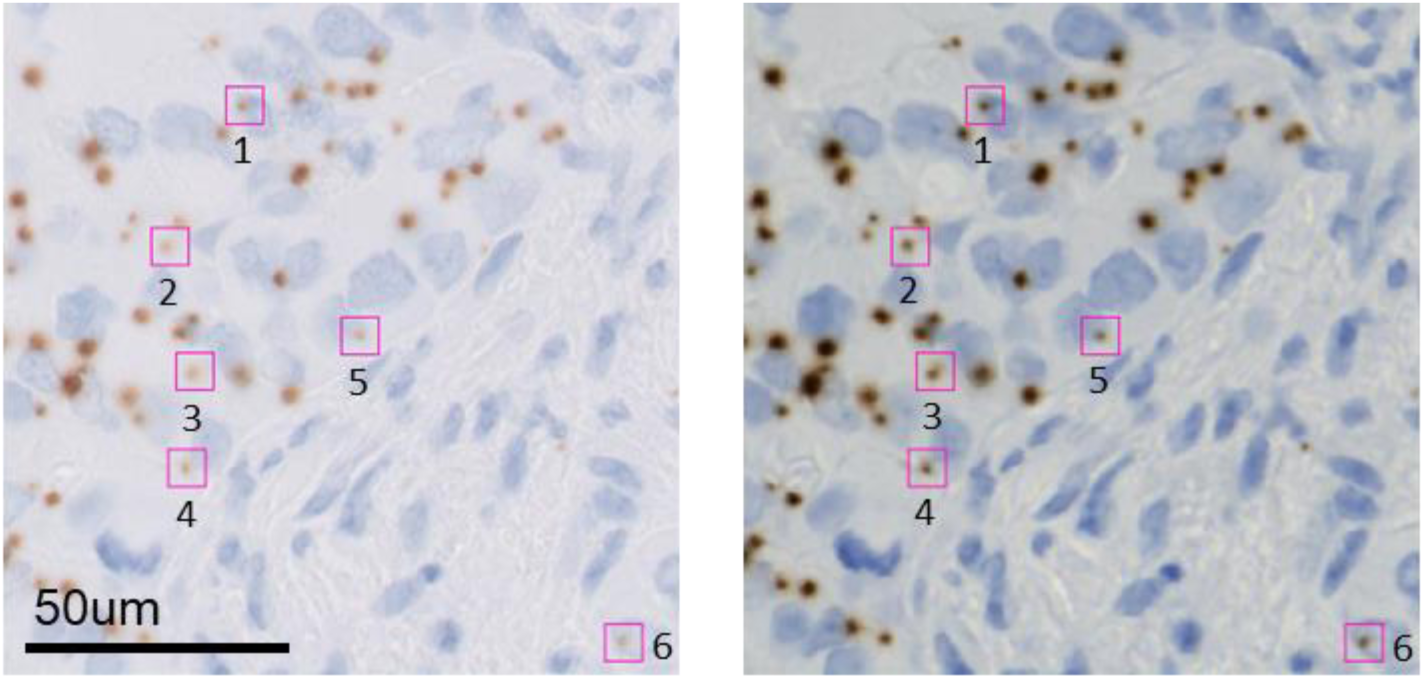
Comparison between single plane and extended depth of focus (EDF) for dot detection. Left image represents a single plane field of view from a HER2 qIHC stained breast cancer section scanned on the Philips Ultra Fast scanner. The magenta boxes highlight six faint qIHC dots that might be considered as staining artifacts instead of real dots. Right image represents the same slide area scanned using the extended depth focus option on the Zeiss Axioscan.Z1 with software-based flattening of five focus planes, dots are clearly observed.

**Figure S4:**
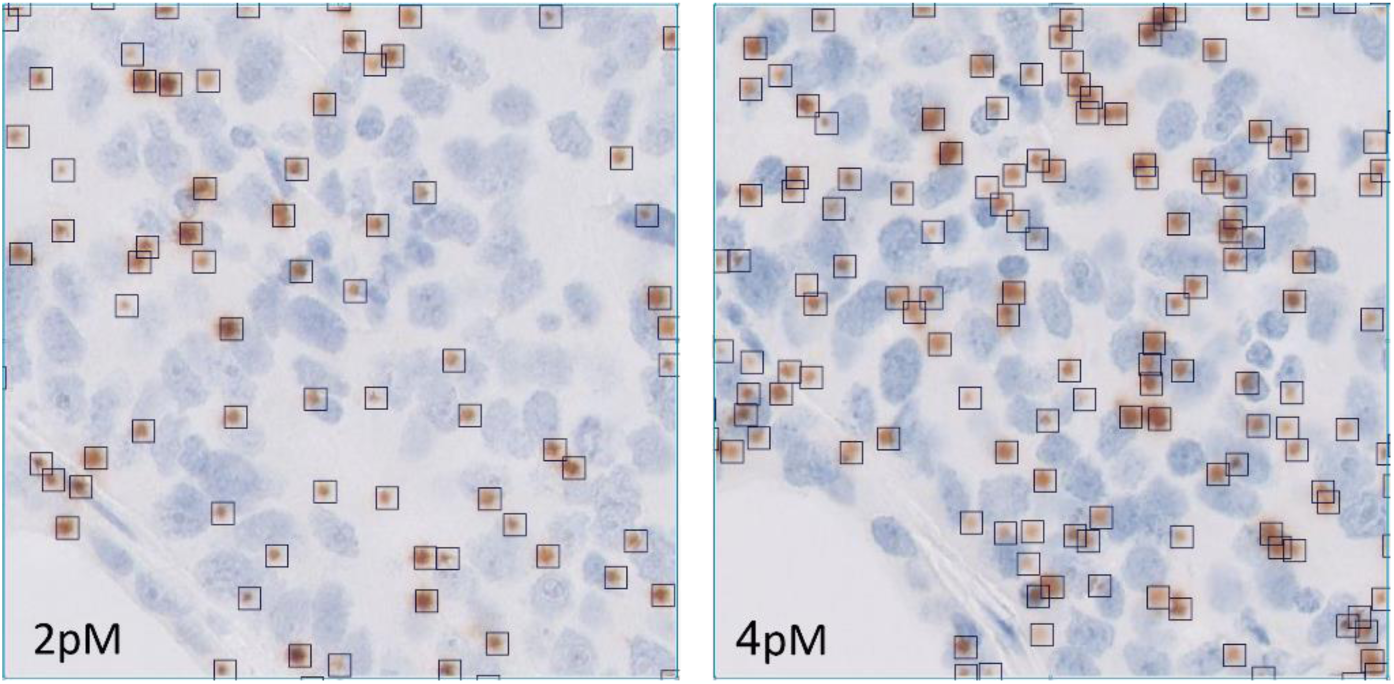
Consecutive tissue sections stained by qIHC using different concentrations of the HRP-conjugated polymer. Left section was stained with 2 pM, while the right section was stained with 4 pM. In sections where qIHC dot density becomes relatively high, dot counts are extracted from the lower concentration (2 pM) section to ensure accurate quantification of dots. Dot counts were 69 and 131 for the 2 pM and 4 pM sections, respectively.

**Figure S5.**
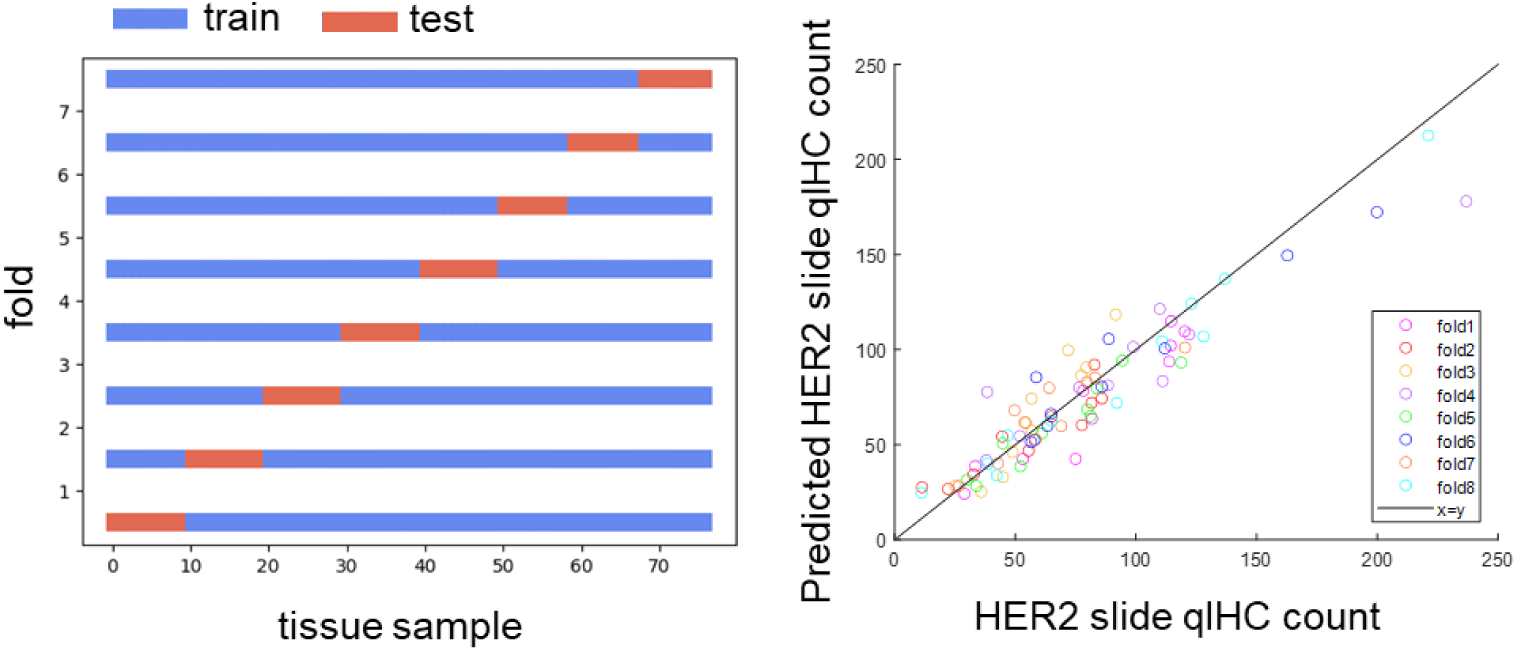
Principle of k-fold validation. Left panel illustrates the principle of k-fold cross validation where 82 breast cancer specimens were subdivided into eight non-overlapping subsets, each containing 72 (or 71) specimens for AI-training, and 10 (or 11) for testing the resulting AI-model. The different colors in the right panel illustrate the predicted HER2 qIHC dots per patch versus the actual HER2 qIHC dots per patch from the eight separate train and test events. The line has been drawn to illustrate the perfect correlation where y=x.

**Figure S6:**
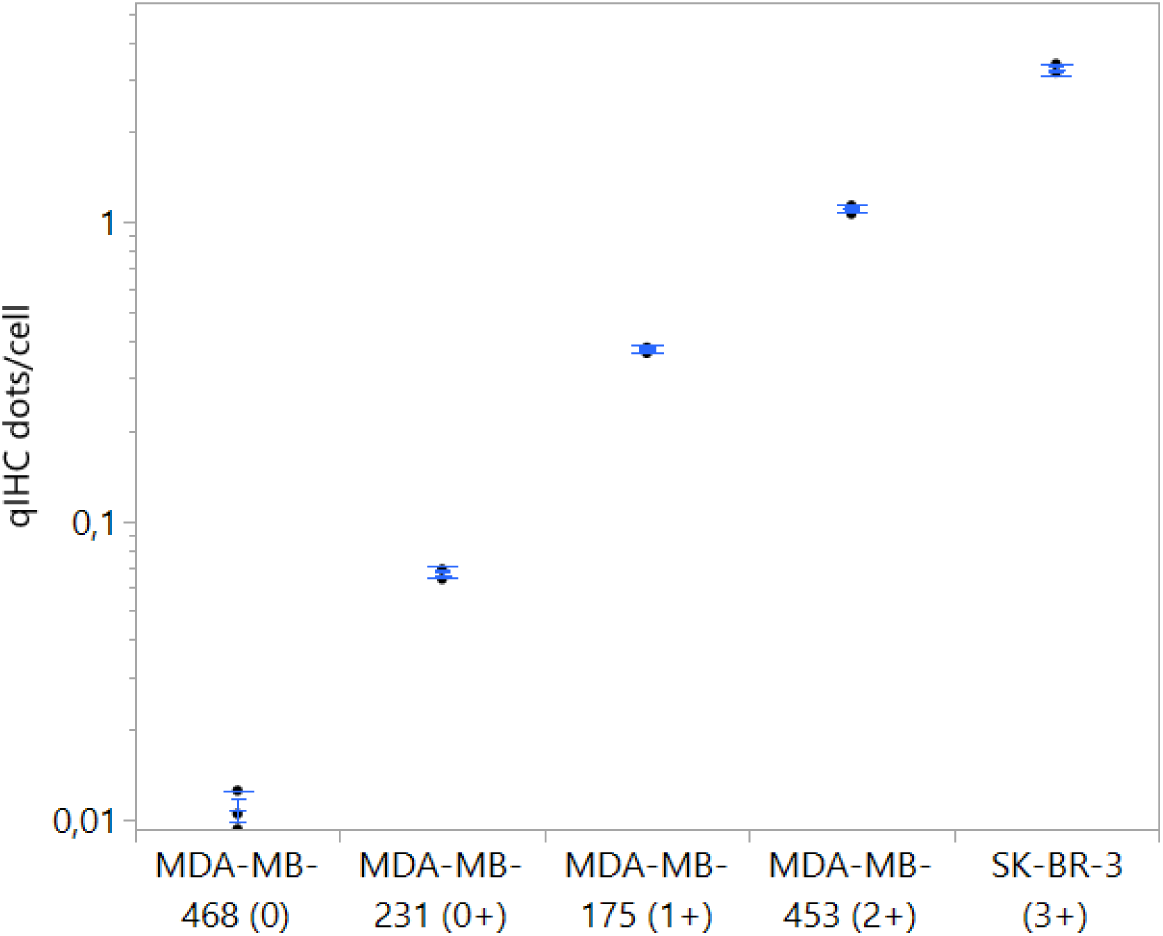
Sensitivity and variability of qIHC in different HER2 expressing cell lines. HER2 qIHC dots/cell in five different breast cancer cell lines using the monoclonal rabbit anti-HER2 antibody (clone DG44) tested in triplicates at a concentration of 1 pM of HRP-conjugated goat anti-rabbit dextran polymer. Bars show the 95% confidence limits of the mean for each cell line.

**Figure S7:**
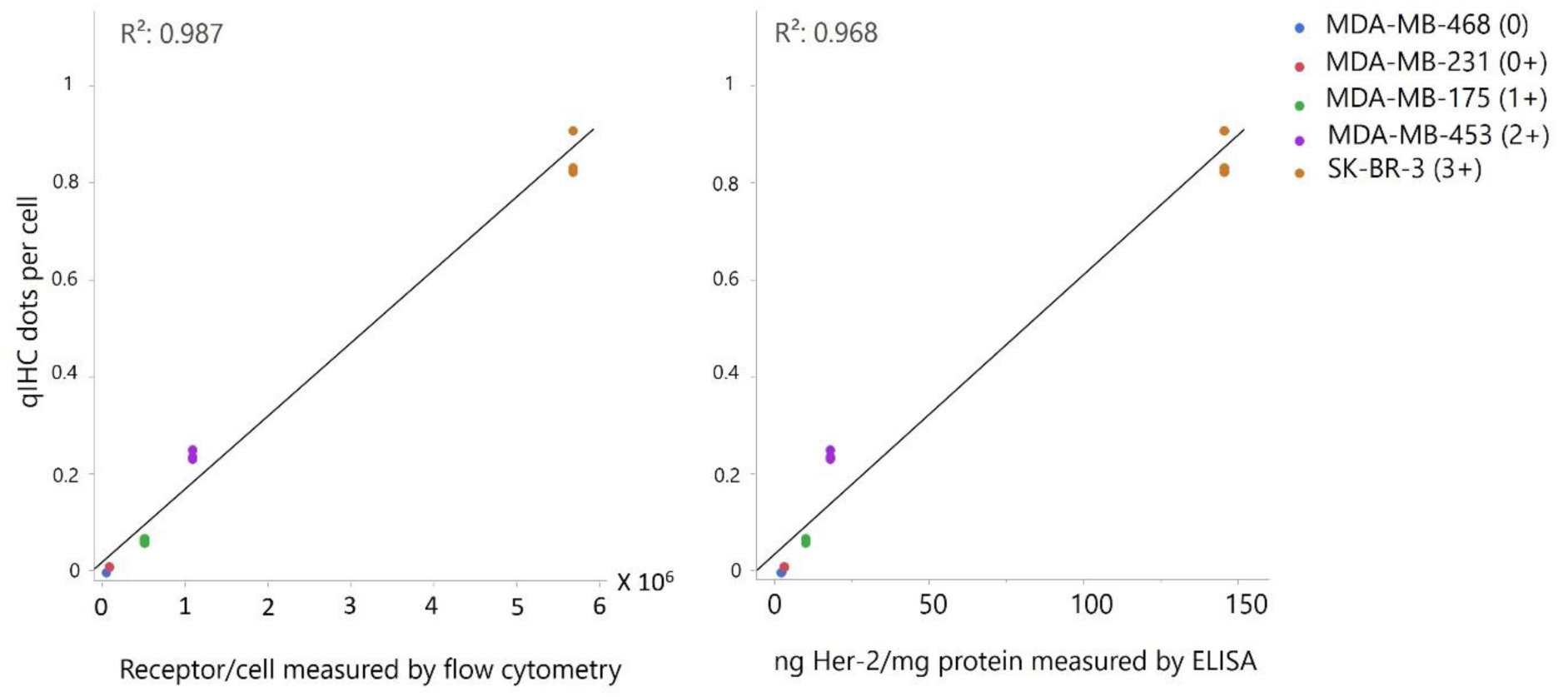
Linearity of qIHC measurement against orthogonal measurements of HER2 in different HER2 expressing cell lines. HER2 qIHC dots/cell in five different breast cancer cell lines using the monoclonal rabbit anti-HER2 antibody (clone DG44) tested in triplicates at a concentration of 0.25 pM of HRP-conjugated goat anti-rabbit dextran polymer. The orthogonal measurement is either an FDA-approval ELISA test for HER2 or flow cytometry. Every measurement is carried out on a sub-portion of the cell line material that was cast into the FFPE block subsequently used for qIHC. Details can be found in Jensen et al., 2017^18^.

